# Super-Resolution Imaging by Dual Iterative Structured Illumination Microscopy

**DOI:** 10.1101/2021.05.12.443720

**Authors:** Anna Löschberger, Yauheni Novikau, Ralf Netz, Marie-Christine Spindler, Ricardo Benavente, Teresa Klein, Markus Sauer, Dr. Ingo Kleppe

**Affiliations:** ZEISS Research Microscopy Solutions, Carl Zeiss Microscopy GmbH, Advanced Development, Carl-Zeiss-Promenade 10, 07745 Jena, Germany; Department of Cell and Developmental Biology, Biocenter, University of Würzburg, Am Hubland, 97074 Würzburg, Germany; Department of Biotechnology and Biophysics, Biocenter, University of Würzburg, Am Hubland, 97074 Würzburg, Germany

**Keywords:** super-resolution microscopy, structured illumination microscopy, 3D multicolor super-resolution imaging

## Abstract

Three-dimensional (3D) multicolor super-resolution imaging in the 50-100 nm range in fixed and living cells remains challenging. We extend the resolution of structured illumination microscopy (SIM) by an improved nonlinear iterative reconstruction algorithm that enables 3D multicolor imaging with improved spatiotemporal resolution at low illumination intensities. We demonstrate the performance of *dual iterative* SIM (*di*SIM) imaging cellular structures in fixed cells including synaptonemal complexes, clathrin coated pits and the actin cytoskeleton with lateral resolutions of 60-100 nm with standard fluorophores. Furthermore, we visualize dendritic spines in 70 µm thick brain slices with an axial resolution < 200 nm. Finally, we image dynamics of the endoplasmatic reticulum and microtubules in living cells with up to 255 frames/s.

## 1. Introduction

Fluorescence microscopy is the most popular imaging method in cell biology because it permits molecule-specific labeling, minimal perturbation and three-dimensional (3D) imaging with single-molecule sensitivity.^[1]^ With the introduction of super-resolution (SR) imaging techniques the diffraction-imposed spatial resolution has been extended by an order of magnitude and enabled the visualization of previously hidden structural details of cells and organelles.^[2,3]^ However, while single-molecule localization microscopy (SMLM) methods can attain virtually molecular resolution insufficient labeling with common fluorescent probes impedes at present the translation of high localization precisions into true molecular resolution.^[4-6]^ Similarly, stimulated emission depletion (STED) microscopy^[2,7]^ and reversible saturable optical fluorescence transitions (RESOLFT) microscopy^[8,9]^ can achieve high spatial resolutions but likewise place extraordinary demands on the photostability and photoswitchability of dyes. Especially in live-cell experiments where lower irradiation intensities and longer wavelengths are key to minimize photodamage^[10]^ resolution expectations have to be winded down. Furthermore, 3D multicolor SMLM, STED and RESOLFT microscopy remain challenging due to the limited number of fluorophores that fulfill the demanding photophysical properties and required expensive machinery as well as extensive expertise.

In this regard, SIM represents a welcome alternative since it enables multicolor imaging with conventional fluorophores and fluorescent proteins at high temporal resolution. In addition, SIM requires substantially lower irradiation intensities and is thus the method of choice for live-cell SR experiments.^[11-13]^ However, SIM still exhibits limitations that restrict its practicability. First, and most importantly, the resolution of SIM is only twice beyond the diffraction limit and the image quality is strongly controlled by the signal-to-noise ratio (SNR) especially in living samples. Second, parts of the sample with strong fluorescence in outer focal areas often generate processing artifacts, which have triggered controversial discussions about data interpretation.^[14-16]^ Finally, even though highly parallelized the number of images required to reconstruct a SR image is still significant. For example, 3D-SIM usually requires 15 images, which is the speed-limiting factor for live-cell experiments.^[17]^

Thus, under practical usage a gap exists between the 10-50 nm resolution provided by SMLM and STED microscopy, and SIM with a resolution of ∼100 nm. A method that enables 3D multicolor SR imaging with 50-100 nm spatial resolution with standard fluorophores and fluorescent proteins in fixed and living cells, and high axial sectioning for tissue imaging combined with a high temporal resolution would close this gap and find widespread applications in biomedical imaging. Here, we extended the resolution of classical SIM by an improved nonlinear iterative reconstruction algorithm. We demonstrate the potential of dual iterative SIM (*di*SIM) by multicolor fluorescence imaging in fixed and living cells, and brain slices with 60-100 nm spatial and millisecond temporal resolution using standard fluorophores and fluorescent proteins under low illumination intensities.

## 2. Results

### 2.1. Improving the Resolution of SIM

In recent years, techniques have been developed to improve the resolution of SIM but often compromises had to be accepted at the expense of possible application fields and the complexity of the microscope. Using objectives with a numerical aperture of 1.7 extends the resolution of SIM towards the sub-100 nm region but is restricted to TIRF illumination^[14,18]^ To achieve higher resolutions nonlinear SIM (NL-SIM) by patterned saturation or fluorescence excitation or patterned depletion of photoswitchable fluorophores has been introduced.^[14,19-21]^ Yet, demanding requirements on the switching properties of fluorophores limit the applicability of NL-SIM for routine life science experiments. Nonlinear SIM also requires significantly more images and thus is in practice limited to 2D imaging.^[14,21-23]^ Alternatively, iterative reconstruction algorithms usually based on Richardson Lucy deconvolution methods have been explored to improve the resolution of SIM.^[18,24,25]^ However, all the methods fell short of expectations and demonstrated that extending the resolution of SIM significantly below 100 nm requires either additional images combined with a limited choice of fluorophores, or the third dimension has to be sacrificed, or even both.^[26]^

To minimize artifacts induced by frequency-shifted signals of out-of-focus background signal total internal reflection fluorescence (TIRF)-SIM^[27]^ and grazing incidence illumination (GI)-SIM^[28]^ have been introduced. Confining the illumination to a small slice close to the coverslip out-of-focus background can be reduced. Due to the reduction of the number of planes that have to be imaged these methods volunteer for live-cell 3D SR-imaging. Other approaches to reduce the out-of-focus background are based on line-scan illumination^[29]^ or 2D illumination patterns.^[30]^ In addition, software algorithms have been used to suppress the background, usually by Fourier filters in the reconstruction process,^[25,31]^ special 2D to 3D deconvolution for 2D data sets^[32]^ and an initial deconvolution (DCV) before the actual SIM processing.^[24]^ A different, but also additional route to reduce artifacts arising from processing of noisy data represents Hessian denoising, which was successfully applied for processing of 2D SIM data with low exposure times.^[18]^ In practice, optimal parameter selection for image reconstruction remains challenging, e.g. resolution enhancement in one part of the image might be accompanied by artifacts in other image regions with more background and noise.^[31]^

Finally, since the number of phase images is linked to the number of spatial frequencies of the illumination pattern 3D SIM acquisition requires two additional phase images per direction. Hence, 2D TIRF-and GI-SIM does not only reduce out-of-focus background signal but likewise enables higher frame rates and thus faster imaging, which is especially important to observe dynamic processes at SR in living cells.^[14,28,33]^ A minor reduction in the number of raw images can be realized by employing a checkerboard type lattice illumination pattern generated by a five beam interference (Lattice-SIM),^[34]^ which only requires 13 phase images.^[22]^ A more significant reduction has been realized by combining the thick slice reconstruction method with stacked 3D acquisition leading to a 3-fold reduction in required images within a z-stack.^[32]^

### 2.2. A Dual Iterative Reconstruction Algorithm

To extend the resolution and speed and minimize artifacts of SIM for 3D imaging we reconsidered iterative reconstruction algorithms.^[35]^ In general, nonlinear algorithms for deconvolution can extend the resolution by reconstructing frequencies outside the support of the optical transfer function (OTF) if combined with a priori boundary conditions such as non-negativity.^[36,37]^ Commonly, linear SIM reconstruction bases on the use of a generalized Wiener filter concept^[17]^ where deconvolution and order combination are processed in a single integrated step such as compact hardware-based solutions combine multiple image manipulations in a single optical element (**Figure 1**). On the other hand, separate consecutive steps offer the advantage to optimize individual steps while possibly reducing interdependencies of the manipulations. To employ effectively nonlinear iterative deconvolution methods, we used two-step processing for SIM reconstructions,^[24,25]^ which enables an improvement in spatial resolution and also a reduction of artifacts related to the Wiener filter. Two-step processing combines advantageously SIM reconstruction including order combination, denoising and frequency suppression filters in the first step with iterative deconvolution in the second step. A point-spread-function (PSF) that is used at the second deconvolution step contains already the effects of digital image manipulations occurring at the first step. Consequently, the reduction of data from 13 phase images into a single SR image in the first step makes the iterative deconvolution more efficient.

**Figure 1.**
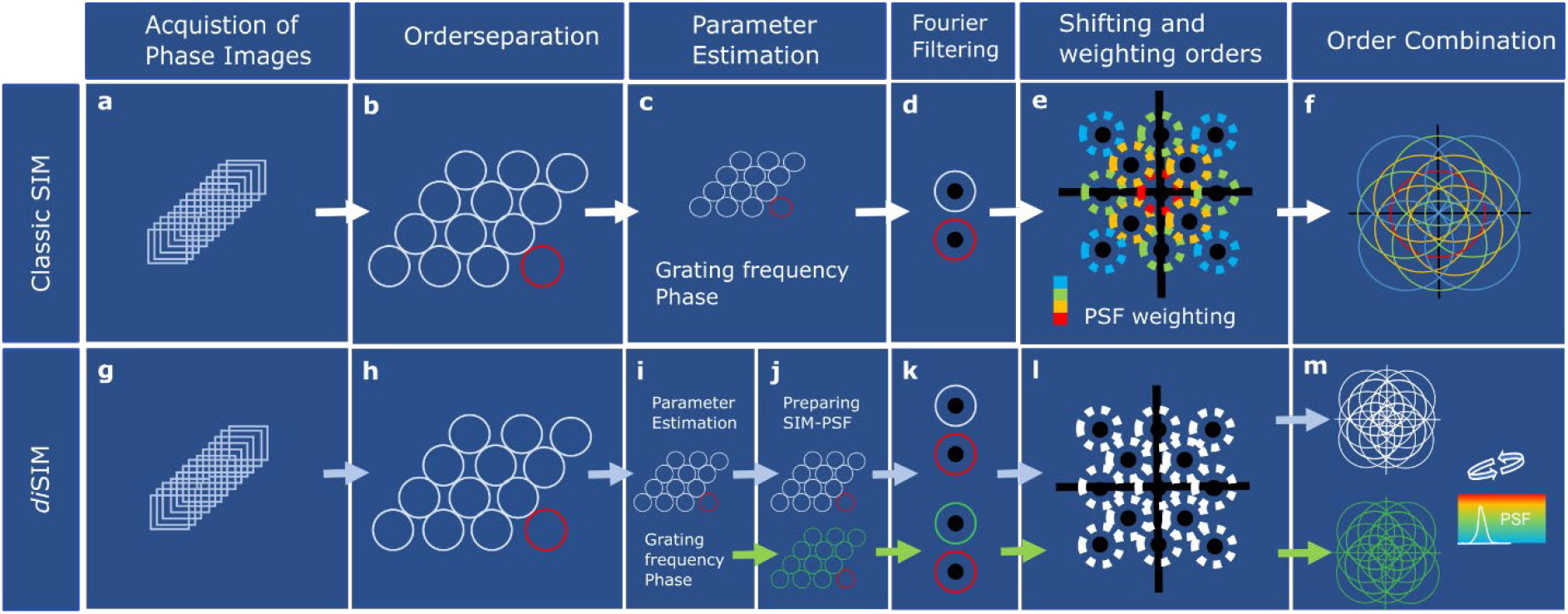
Generalized Wiener reconstruction (Classic SIM) compared to *di*SIM reconstruction. a,g) 13 acquired phase images for Lattice-SIM are first processed to separate orders (frequency bands) in Fourier space corresponding to the demodulation of the Moiré patterns (b,h). c,i) In a second step the fit grating frequency and phase positions are accurately determined from the image data shown in blue. In *di*SIM these parameters are used to generate an image set and start the SIM-PSF generation (green) with the same steps as the data further processed. Potentially the separated orders are additionally unmixed in *z* in case of 2D acquisitions or 3D thick slice reconstructions.^[32]^ d,k) To reduce artifacts and enhance optical sectioning, Fourier filters are applied. Out of focus background is suppressed by applying a high-pass filter on the 0^th^ order (red circle) while the notch filters on the other orders suppress the grating frequencies. e) The next step shifts and places the orders to the appropriate places in frequency space and weights these orders according to the illumination and optical transfer function (OTF) (colors) to boost high frequency information which otherwise would drown in the noise when combining all orders in the next step. f) The combination of the orders into the SR image for Classical SIM is done by a generalized Wiener filter. l,m) In *di*SIM orders are only weighted by the illumination strength but not weighted according to the OTF and combined by simple summation. However, the PSF for the subsequent iterative deconvolution, which is the final step, is generated in the same way (green arrows) as used for image reconstruction by processing a point response with the exact same parameters, extracted from the data.

Like in optics, the elements in image formation, including unwanted properties such as aberrations, must be reflected in the PSF to avoid artifacts, and in most cases, to enable conversion of the algorithm. Since iterative deconvolution critically depends on the correct underlying model, a PSF that contains all effects, also of the digital image manipulations is key for accurate image reconstruction. Thus, the incorporation of all initial processing steps into the effective SIM-PSF for image formation is a crucial point. In purely hardware-based imaging techniques this would be addressed by employing an experimental PSF, which contains all systematic aberrations. Analogously, we developed *dual iterative* SIM (*di*SIM), a processing method that uses dual reconstruction by preparing the data and a digital experimental PSF in the same way before entering the second step of deconvolution (Figure 1).

### 2.3. *di*SIM Imaging with Improved Spatial Resolution

Since the achievable resolution and quality of reconstructions are generally sample dependent, we imaged different technical and biological samples with known underlying structure to validate the method. To assess the performance of *di*SIM we first used 60 nm DNA origami structures,^[38]^ and a re-usable stable fluorescence slide with linear patterns (Argo-SIM slide) (**Figure 2**, Figure S1, Supporting Information). The peak-to-peak distance of 60.2±9.4 nm (s.d.) determined for the DNA origamis as well as the distances resolved on the Argo-slides demonstrate unequivocally that *di*SIM provides a lateral resolution ∼60 nm. To demonstrate the potential of *di*SIM for SR imaging of multiprotein complexes we selected the synaptonemal complex (SC) as a biologically relevant and complex 3D reference structure (**Figure 3**, Figure S2, Figure S3, Supporting Information).

**Figure 2.**
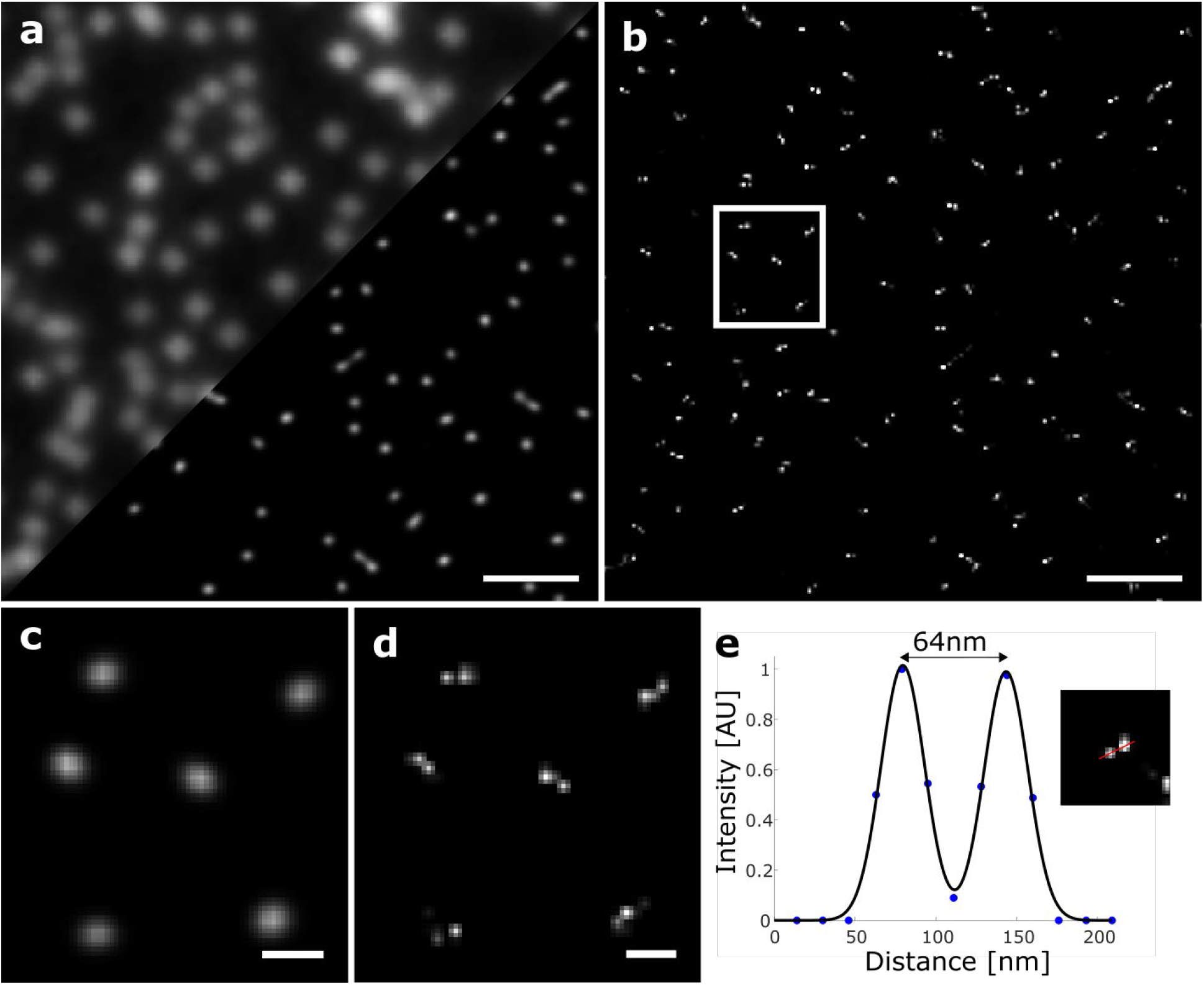
*di*SIM enables 60 nm spatial resolution. a) Fluorescence images of 60 nm origami structures (upper left wide-field, lower right standard linear SIM reconstruction). b) *di*SIM image of 60 nm origami reconstructed with the iterative reconstruction algorithm with a digital experimental PSF. c,d) Expanded views of the region marked by a white rectangle in (b) comparing standard SIM (c) with *di*SIM (d). e) A Gaussian fit of the intensity profile of a selected 60 nm origami delivered a peak-to-peak distance of 64 nm. The peak-to-peak distance of 25 randomly chosen origamis was determined to 60.2±9.4 nm (s.d.). Scale bars, a,b) 1 µm, c,d) 200 nm.

**Figure 3.**
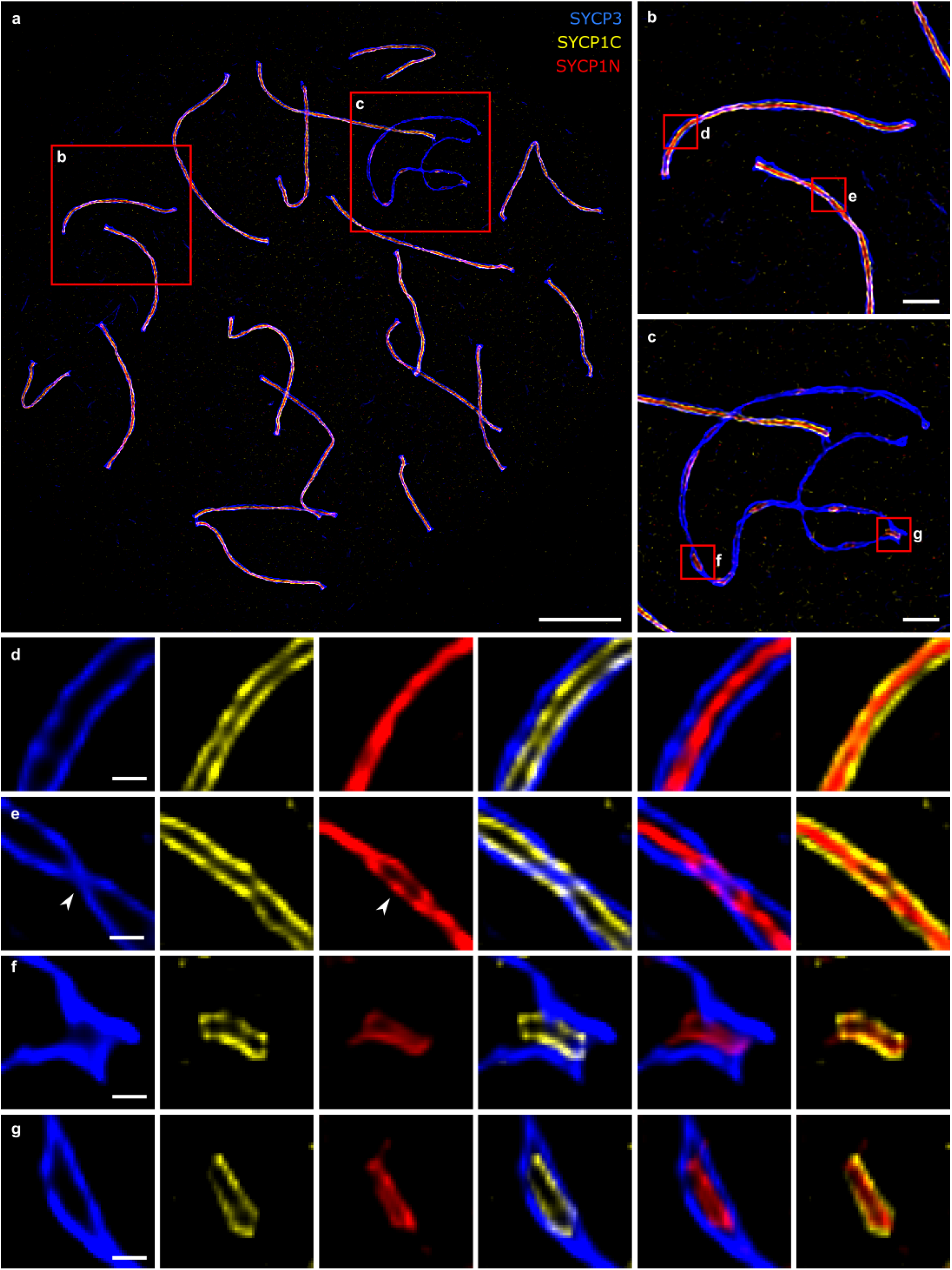
Three-color *di*SIM images of murine spreadings of the SC demonstrate its superior spatial resolution. a) Morphologically, the SC consists of two lateral elements that are connected with a central element via a multitude of transverse filaments. Lateral element protein SYCP3 was labeled with a SeTau647 coupled secondary antibody (blue), the C-terminus of the transverse filament protein SYCP1 with Alexa Fluor 488 antibody (yellow), and the SYCP1 N-terminus with Alexa Fluor 568 antibody (red). b,c) Magnified images of the red rectangles in (a) show SCs that are formed between autosomes (b) and stretches of the SC along the XY pair (c). d-g) Magnified views of red rectangles in (b,c). Twisting of the complex around its own axis results in the exposure of frontal (d) and lateral view (e) sections. Protein distribution analysis of SYCP3 and SYCP1 based on *d*STORM data of frontal view SC sections revealed bimodal distributions of SYCP3 and SYCP1-C with peak-to-peak distances of roughly 220 nm and 148 nm, respectively, as well as a monomodal signal distribution of roughly 40 nm for SYCP1-N.^[40]^ Accordingly, *di*SIM visualizes two strands of SYCP3, two strands of SYCP1-C and a single strand of SYCP1-N in frontal views (d). Most importantly, the signals of the three proteins are resolved separately. Lateral view sections, which are equivalent to sites where the SYCP3 signal twists (e, arrowhead), *di*SIM resolves a splitting of the SYCP1-N signals (Figure S2, Supporting Information).^[40,42]^ Furthermore, *di*SIM resolves the SC proteins individually both at the pseudo-autosomal region of the XY pair (f) as well as at the unsynapsed regions of the sex chromosomes (g). Scale bars, a) 5 µm, b,c) 1 µm, d-g) 250 nm.

The SC is a hallmark structure of the meiotic nucleus where it facilitates recombination and haploidization in sex cells across species.^[39]^ The nanoscale dimensions of the SC have been well characterized by different SR techniques including *d*STORM^[40]^ and combinations of SR and expansion microscopy.^[41,42]^ However, while SMLM methods are restricted to one-or two-color imaging using suited fluorophores in photoswitching buffer, three-color *di*SIM with standard fluorophores resolves the organization of the SC with a resolution previously only achieved by three-color SIM of ∼3-fold expanded SCs (Figure 3, Figures S2, S3, Supporting Information).^[42]^ This allowed us to resolve a splitting in the signal of the N-terminally labeled transverse filament protein SYCP1 (SYCP1-N) in lateral views of the SC (Figure 3e). To demonstrate the high spatial resolution of *di*SIM we analyzed the distance between the SYCP1-N signals for selected regions close to the twisting points and determined values between 48 and 59 nm (Figure S2, Supporting Information). In addition, *di*SIM unravels the bifurcation and fraying of the lateral element protein signal (SYCP3), which coincides with the fraying of the chromosome axis (Figure S3, Supporting Information).^[42]^ Furthermore, *di*SIM resolves the multistrandedness of the SC’s lateral elements (Figure 3), and the protein distributions at the pseudo-autosomal region of the XY pair and at unsynapsed regions of the sex chromosomes (Figures 3f, 3g). These results demonstrate that multicolor *di*SIM is ideally suited to resolve fine structural details of the organization of multiprotein complexes with a resolution approaching 50 nm in efficiently labeled samples with high SNR.

Next, we used clathrin-coated pits (CCPs) as cellular reference structures for SR imaging.^[14,43,44]^ CCPs are responsible for endocytosis of several plasma membrane receptors and their cargos. We used immunolabeling and saw that microtubules and CCPs were resolved as homogeneously labeled filaments and hollow rings with diameters varying from 83 nm to 163 nm and an average diameter of 122.4 ± 19.5 nm (s.d.) (Figure S4, Supporting Information). The smaller CCP population in *di*SIM images provides corroborating evidence that *di*SIM achieves a spatial resolution far below 100 nm also for imaging of intracellular structures (Figure S4, Supporting Information).

Finally, we selected the nuclear pore complex (NPC) as a challenging SR imaging reference structure to reveal the limits of *di*SIM. NPCs are wheel-shaped, eightfold symmetrical cylindrical protein assemblies with a ∼120 nm diameter core structure surrounding the central channel with a diameter of ∼50 nm.^[45]^ It was shown that immunolabeling and *d*STORM in combination with particle averaging enables the visualization of the eightfold symmetry and determination of the diameters of the core structure and the central channel with a spatial resolution of ∼20 nm.^[46,47]^ *di*SIM images of isolated *Xenopus laevis* oocyte nuclear envelopes immunostained for gp210, an integral membrane protein of the NPC, reveal the resolution limits of *di*SIM. They show the wheel-shaped arrangement of the NPCs (Figure S5, Supporting Information) but cannot resolve details of the molecular organization nor the central channel of NPCs with a diameter of ∼40 nm as has been shown by *d*STORM.^[46]^

### 2.4. 3D Imaging and Optical Sectioning Performance of *di*SIM outperforms classical SIM

The rejection of out of focus light is not only key for imaging deeper into scattering tissue but also most relevant for 3D imaging in biology. Even thin samples such as cultured cells can be challenging if densely labeled. Phalloidin staining of the actin cytoskeleton is such a typical example, where delicate structures can easily be drowned in the background arising from bright structures in outer focal areas (**Figure 4a**, Figure S6, Supporting Information). The background often results in significant artifacts in conventional SIM reconstructions and can hamper scientific interpretation of the data. The improved axial sectioning of *di*SIM completely removes these artifacts (Figure 4a) and allowed us to cut thinner optical sections than conventional SIM (Figures 4b, 4c) with a sectioning capability of < 200 nm (Figure 4b). In contrast to GI-SIM with a 1 µm thin light-sheet and TIRF-SIM with ∼100 nm sectioning *di*SIM is independent of the axial position relative to the coverslip and stackable for 3D volume imaging. Fourier ring correlation analysis^[48]^ demonstrates that *di*SIM achieves a substantially improved lateral resolution of ∼ 50 nm also for more complex cellular structures (Figure S7, Supporting Information).

**Figure 4.**
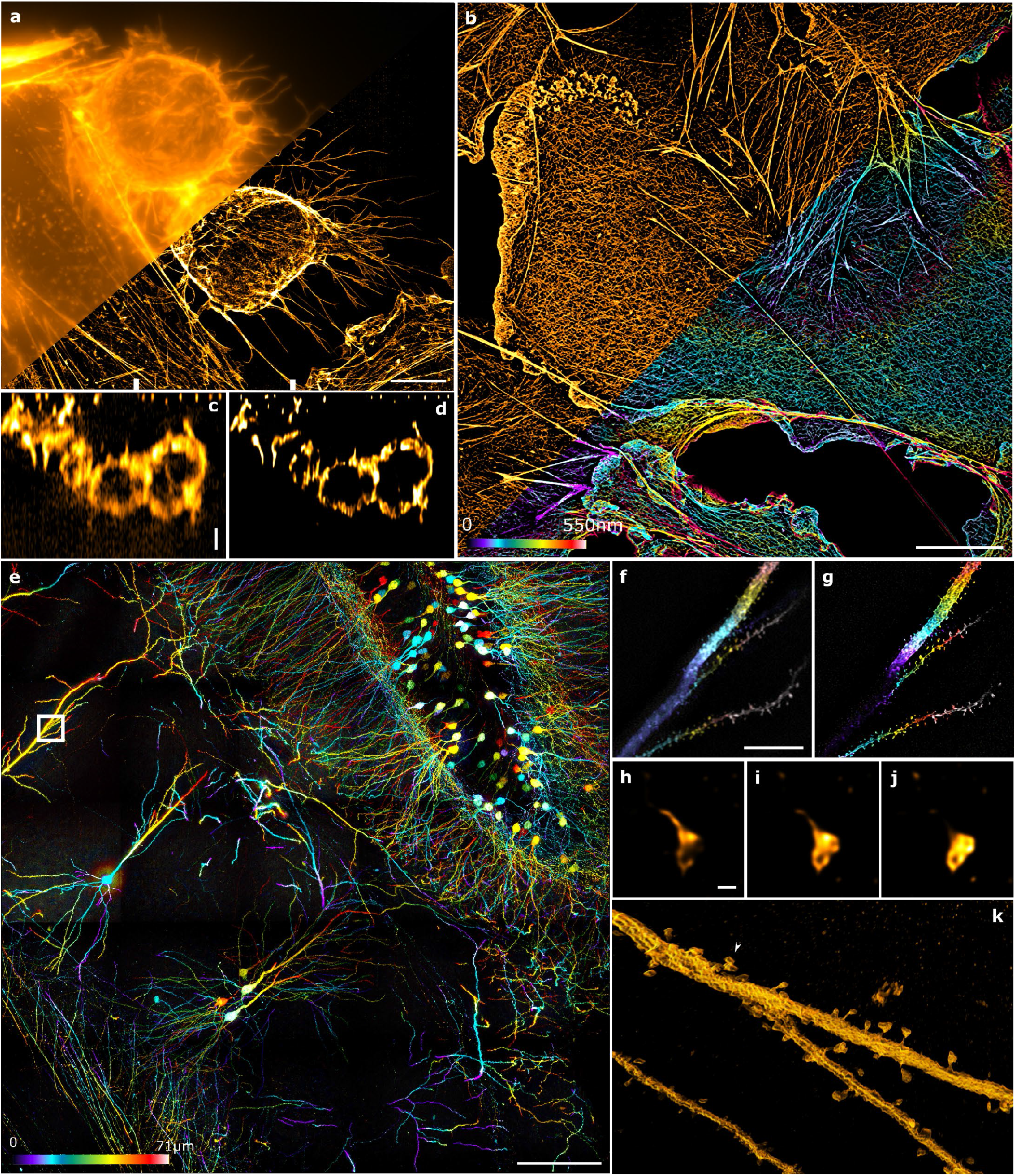
Axial sectioning capabilities of *di*SIM in densely labeled and scattering thick samples. a) Fluorescence image (maximum intensity projections) of the actin cytoskeleton of a U2OS cells stained with Alexa Fluor 568-phalloidin. The upper left part shows a conventional wide-field image and the lower right part a *di*SIM image. b) 3D *di*SIM image of the actin cytoskeleton of a COS-7 cells stained with Alexa Fluor 488 phalloidin. The upper left part shows a 2D maximum intensity projection and the lower right part the 3D color-coded image. c,d) Corresponding *xz* projections of the images shown in (a) with standard SIM processing (c) and *di*SIM (d), *x*-axis region as indicated by the white marks at the bottom of (a). e) 3D *di*SIM image of a 71 µm thick hippocampal slice of a mouse brain of a Thy1-GFP line imaged in 5×5 tiles using a 25x magnification objective and employing a 1D stripe pattern (SIM-Apotome mode). f,g) Color coded maximum intensity projections of the white square in (e) for a reduced *z*-range (0 – 5.7 µm) using conventional SIM (f) and *di*SIM reconstruction (g). h-j) Cup of dendritic spine of a cortical neuron imaged in the same brain slice with higher magnification (63x) by *di*SIM. Images were taken every 110 nm in axial direction (h to j). k) 3D rendering of the *di*SIM reconstruction of a cortical dendritic branch point reveals the cup shaped 3D morphology of the spines. White arrow indicates the spine as shown in (h-j) for comparison of scale. Scale bars, a,b) 10 µm, c,d) 2 µm, e) 100 µm, f,g) 5 µm, h-j) 200 nm.

To explore the use of *di*SIM for tissue imaging, we imaged brain slices of a Thy1-GFP mouse (Figure 4e-4k). Here, we combined 1D SIM stripe patterns (SIM-Apotome mode) for low NA magnifications (Figure 4e-4g) and lattice patterns for high magnification image acquisitions (Figure 4h-4k). The overview image of the hippocampus (Figure 4e) resolves fine details of spines through the entire depth of the brain slice of 71 µm and demonstrates the superior sectioning of *di*SIM compared to conventional SIM.

Recently, dimensional reduction and machine learning has been applied to large data sets to characterize spine morphology based on SIM in cultured neurons demonstrating equivalent results to reconstructions from EM data.^[49]^ Equally, *di*SIM reconstructs fine spine morphology within scattering brain slices at axial depths of several micrometers (Figure 4f-4g). Therefore, we reason that *di*SIM in combination with refined methods of artificial intelligence can pave the way to study physiological connectivity in tissue and potentially *in-vivo* with unprecedented precision.

### 2.5. *di*SIM Enables Live Cell Imaging with High Spatiotemporal Resolution

To demonstrate the performance of *di*SIM for live cell SR imaging we expressed Calreticulin-tdTomato, a highly conserved chaperone protein which resides primarily in the endoplasmic reticulum (ER), and the microtubule binding protein EMTB-3xGFP for monitoring of ER and tubulin dynamics in COS-7 cells (**Figure 5**). To capture fast dynamics without sequential imaging of different axial planes and 3D image stack reconstruction, we extended the *di*SIM concept by combining it with a deconvolution step that generates 3D from 2D data sets (Figure 1).^[32]^ Resolving fine structural details of dense tubular matrices in the peripheral ER remains very demanding because it requires nanoscale resolution on milliseconds timescale.^[49]^ Compared to conventional 2D SIM our new *di*SIM 3D reconstruction recorded with a frame rate of 255 Hz produced sharper images and revealed more tubular details (Figure 5a-5c).

**Figure 5.**
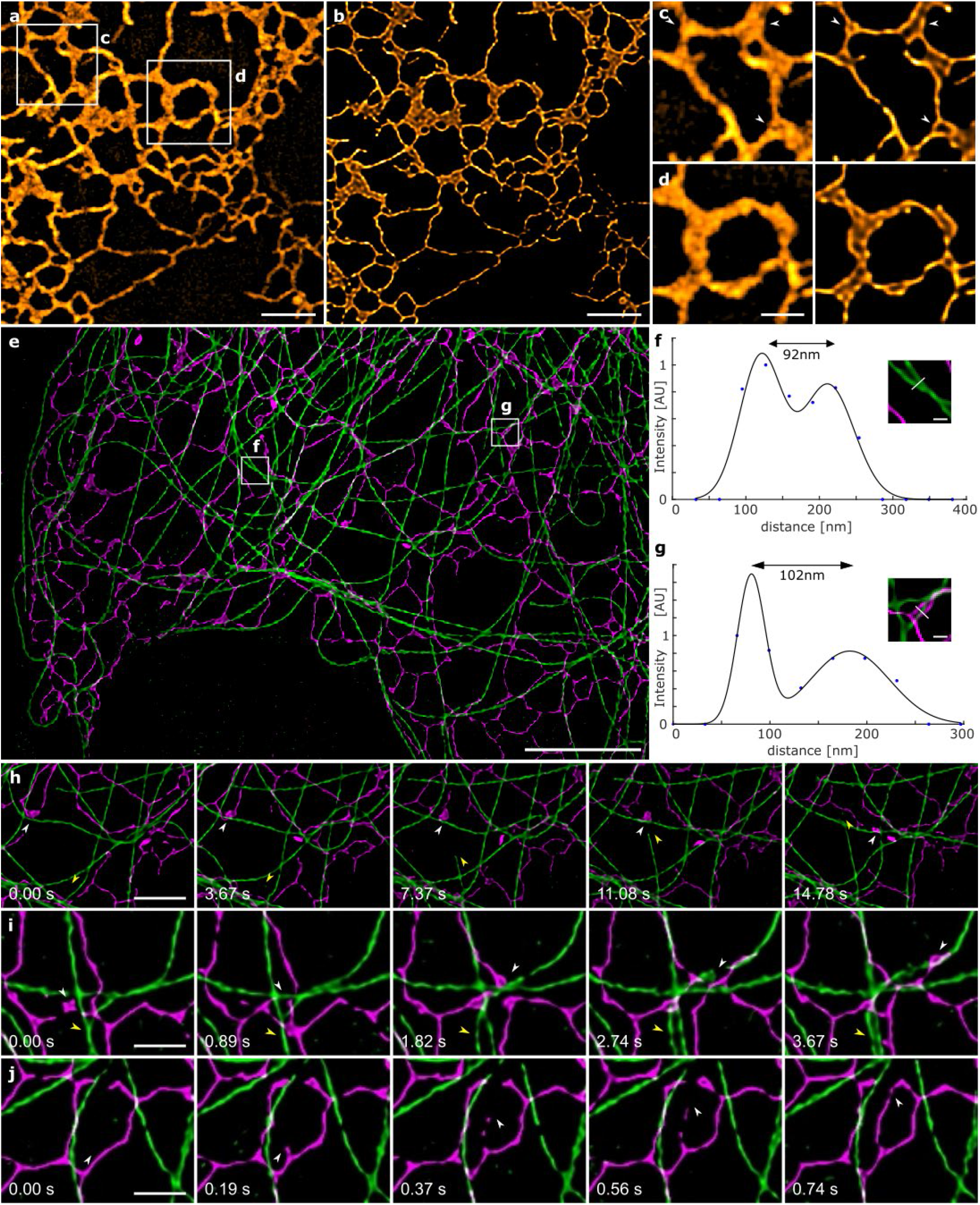
Imaging live-cell dynamics by *di*SIM. a,b) COS-7 cells were transfected with Calreticulin-tdTomato and imaged with a frame rate of 255 Hz by SIM (a) and *di*SIM (b). c,d) expanded views of the white rectangles (a). *di*SIM processing (right) reveals more tubular details than conventional SIM processing (left). For better visibility gamma had been adjusted to 0.45 in the *di*SIM images. e) Simultaneous two-color imaging of Calreticulin-tdTomato and EMTB-3xGFP at a framerate of 81.6 Hz shows highly resolved structures of microtubules (f) and ER (g). h-j) Snapshots taken at different times. h) Every 100^th^, i) every 25^th^, and j) every 5^th^ time point of a time series (Movies S1-S3, Supporting Information). Yellow arrows point out growing (h) and splitting (i) microtubules. White arrows illustrate different dynamics of ER, containing hitchhiking events on microtubules (h,i) and *de novo* budding of ER (j). Scale bars, a,b) 2.5 µm, c,d) 1 µm, e) 5 µm, f,g) 250 nm, h) 2 µm, i,j) 1 µm.

In addition to the high temporal resolution provided by Lattice-SIM, the sectioning capabilities of *di*SIM compared to conventional SIM enabled us to increase the detection of tubular structures in the usually more sheet like looking dense tubular matrices of the peripheral ER.^[28]^

For simultaneous two-color imaging of ER and microtubules we decreased the frame rate to 81.6 Hz to increase the SNR and reduce photobleaching. *di*SIM resolved easily distances of 92 nm and 102 nm between adjacent microtubules and tubules of the ER, respectively (Figure 5f, 5g). Furthermore, *di*SIM revealed several well-known types of movements happening at different timescales including the growth of microtubules, ER hitchhiking and ER budding (Figure 5h-5j, Movies S1-S3, Supporting Information).^[33]^ These data highlight the performance of Lattice-SIM with *di*SIM data processing for 2D two-color live-cell SR imaging with millisecond temporal and sub-100 nm spatial resolution.

## 3. Discussion

Limitations to the *di*SIM method arise from iterative deconvolution in the same way as for other imaging modalities such as widefield, confocal, light-sheet or lattice light-sheet microscopy. These limitations are usually seen by ringing artifacts or spurious structures due to noise propagation into the final image. However, these effects are usually taken care of by using appropriate regularizers and ensuring that the algorithm converges. We applied accelerated maximum likelihood algorithms of iterative conjugate gradient deconvolution^[50]^ using Poisson likelihood, as a maximization target, which delivered robust results also for lower light levels. However, the noise statistics is only approximated by a Poissonian due to the first processing steps, which modifies the underlying statistics slightly. This leaves room for future improvements by adapting the conjugate gradient deconvolution with a more accurate likelihood. In high SNR cases with small amount of background, the basic non-negativity constraint was already good enough to generate high-quality images. In all other cases, we improved results by applying first-order derivative regularizations (Good’s roughness).^[51]^ We typically used 15-20 iterations and, in some cases, up to 40 iterations, which is comparable to widefield deconvolution.

A further restriction of the iterative deconvolution method is the dependency of the final resolution of the image on the sample itself.^[35]^ This is also the reason why we demonstrated the applicability of the method on various technical and biological structures which are known from the literature using other super resolution methods to show the method’s robustness for biological relevant structures. The resolution enhancement we report, from SIM (100 nm -130 nm) to *di*SIM (60 – 90 nm) compares well to numbers being achieved with iterative deconvolution in other imaging modalities. For example, deconvolution of widefield fluorescence images improves the lateral resolution from about 200 – 300 nm to 180 – 250 nm, and confocal fluorescence images can resolve well below even 140 nm using iterative deconvolution, while the best conventional confocal resolution is known to be 180 – 250 nm.^[52-54]^

## 4. Conclusions

*di*SIM improves the resolution of classical SIM and enables multicolor fluorescence imaging with 60-100 nm spatial and millisecond temporal resolution using standard fluorophores and fluorescent proteins. Furthermore, we have demonstrated the sectioning capabilities of *di*SIM emphasizing its advantageous use for SR imaging in tissue. We believe that *di*SIM volunteers as an easy to use and reliable SR imaging method that closes the resolution gap between the 100 nm range provided by standard SIM and the <50 nm range provided by more sophisticated and less live-cell suited methods such as SMLM and STED microscopy. Potential applications include multicolor live-cell studies that require superior spatiotemporal resolution but also multicolor 3D SR-imaging of expanded samples with molecular resolution.^[6,55]^

## 5. Experimental Section/Methods

### Ethics statement

The regulatory agency of the city of Wuerzburg approved the animal housing, breeding and experimental protocols (Reference 821– 8760.00–10/90 approved 05.06.1992; according to §11/1 No. 1 of the German Animal Welfare Act). The animal care and experimental protocols were conducted according to the guidelines provided by the German Animal Welfare Act to insure careful, consistent and ethical handling of mice (German Ministry of Agriculture, Health and Economic Cooperation).

### Reagents

Bovine Serum Albumin (BSA, A2153, Sigma), Di-sodium hydrogen phosphate dehydrate (A3567, Applichem), 1,4-Dithiothreitol (DTT, A1101, Applichem), ethylenediamine-tetraacetic acid (EDTA, E1644, Sigma), normal goat serum (50197Z, Thermo Fisher), paraformaldehyde (31628.02, Serva), PBS (P5493, Sigma), poly-D-lysine hydrobromide (P6407, Sigma), potassium chloride (A2939, Applichem), potassium di-hydrogen phosphate (A1043, Applichem), sodium chloride (NaCl, S7653, Sigma), sucrose (S0389, Sigma), Tris base (T6066, Sigma), trisodium citrate dehydrate (S1804, Sigma), Tween-20 (28320, Sigma), Triton X-100 Surfactant (T8787, Sigma).

### Meiotic nuclear spreadings

For meiotic nuclear spreadings, male wildtype C57.6J/Bl6 between the ages of 14-days to 12 weeks were sacrificed using CO_2_ and subsequent cervical dislocation. For nuclear spreadings the testes were resected and decapsulated in PBS to extract the seminiferous tubules.^[56]^ The tubules were transferred to hypotonic buffer (30 mM Tris/HCl, 17 mM sodium citrate, 5 mM EDTA, 50 mM sucrose, 5 mM DTT) and incubated for 60 minutes. Individual seminiferous tubules were then transferred to sterile 100 mM sucrose solution on a slide. The tubules were mechanically disrupted in the sucrose solution using tweezers. In the next step, the testes cells were released into the solution by resuspension with a 10 µl pipette. A drop of 1% paraformaldehyde was collected on the edge of a poly-lysine slide that was previously immersed in 1% PFA. 20 µl of the testis cell solution were transferred into the drop of formaldehyde and spread evenly across the entire slide. The nuclear spreadings were incubated in a humidified chamber for 2 hours and dried for at least 16 hours with the lid of the chamber left ajar. Dried down slides were either used immediately or stored at -80°C in aluminum foil and thawed prior to use. Care was taken, that thawed slides were brought to room temperature prior to unwrapping from the aluminum foil to avoid condensation on the sample.

### Antibodies

Antibodies for the detection of the N-terminus of guinea pig anti-SYCP1 and the C-terminus of rabbit anti-SYCP1 (aa 922-997) were generated by Seqlab.^[57-59]^ The resulting polyclonal antibodies were purified by affinity chromatography. Mouse anti-SYCP3 (ab97672; full length protein) was purchased from Abcam. Table 1 and Figure S9 (Supporting Information) give an overview of the primary antibodies used for the immunolocalization of synaptonemal complex proteins on nuclear spreadings. SeTau647NHS (K9-4149) was purchased from SETA Biomedicals and conjugated to F(ab’)2 of goat anti-mouse IgG (SA-10225) of ThermoFisher. Al568 goat anti-rabbit IgG (H+L) highly cross-adsorbed (A-11036); Al488 F(ab’)2 goat anti-mouse IgG (H+L) cross-adsorbed (A-11017); Al568 F(ab’)2 anti-mouse IgG (H+L) cross-adsorbed (A-11019); Al488 F(ab’)2 goat anti-rabbit IgG (H+L) cross-adsorbed (A-11070); Al647 goat anti-guinea pig IgG (H+L) highly cross-adsorbed (A-21450); Al568 goat anti-guinea pig IgG (H+L) highly cross-adsorbed (A-11075) were purchased from ThermoFisher. Al488 F(ab’)2 goat anti-guinea pig affinity purified (106-546-003) was purchased from Dianova.

### Immunolocalization on nuclear spreadings

Thawed nuclear spreadings were washed with PBS and incubated with 500 µl 10% NGS for 1 hour at room temperature in a humidified chamber to block unspecific epitopes. The blocking solution was drained and the spreadings were incubated with 150 µl of the primary antibody diluted in PBT (1 hour; humidified chamber; for dilutions see Table above). After washing with PBS, the spreadings were blocked for another 30 minutes, before they were incubated with 150 µl of the secondary antibody diluted in PBT (1:200) for 30 minutes at room temperature in a humidified chamber. For dual and triple immunolocalizations, immunostaining was performed sequentially. Spreadings were embedded in ProLong Glass Antifade mountant (P43984; ThermoFisher).

### Phalloidin labeling of COS-7 cells

Cells were seeded on coverslips (thickness of 170 µm) and cultivated for two days. They were rinsed in prewarmed (37°C) PBS. Fixation followed similar to the protocol of Small *et al*.^[60]^ Cell were permeabilized for 1-2 min in 0.3% glutaraldehyde and 0.25% Triton X-100 in PBS. Then they were fixed for 10 min in 2% glutaraldehyde in PBS. After two short washing steps in PBS a 7 min incubation in 0.1% NaBH4/PBS followed for background quenching. They were washed 2 times and incubated in phalloidin Alexa Fluor 488 in PBS over night at 4°C and stored there until imaging. For imaging the phalloidin solution was replaced with PBS.

### Immunostaining of clathrin coated pits in COS-7 cells

Cos7 cells were seeded on 18 mm 1.5 H coverslips (Carl Roth, LH23.1) which were placed in a 12-well plate (50 000 cells per well). The next day cells were rinsed with warm cytoskeleton buffer (CB: 10 mM MES pH 6.1, 138 mM KCl, 2 mM EGTA, 320 mM sucrose, 3 mM MgCl2), fixed with warm 4% formaldehyde (Merck, F8775) in CB for 10 minutes, washed twice with PBS and once with 100 mM glycine in PBS for 5 minutes. After two minutes permeabilization with 0.1% Triton X-100 (ThermoFisher, 28314) in PBS and washing three times with PBS, cells were blocked with 5% BSA (Merck, A3983) in PBS for 30 minutes. Mouse anti-clathrin (abcam, ab2731) and rabbit anti-alpha tubulin (abcam, ab18251) primary antibodies were diluted 1:200 in 5% BSA in PBS, resulting in a concentration of 30 µg/ml and 5 µg/ml, respectively. After 60 minutes incubation at room temperature with the primary antibodies cells were rinsed with PBS and washed twice for 5 minutes with 0.1% Tween-20 (ThermoFisher, 28320) in PBS. As secondary antibodies goat anti-rabbit IgG Alexa Fluor 568 (ThermoFisher, A-11036) and goat anti-mouse F(ab’)2 fragment Alexa Fluor 488 (ThermoFisher, A-11017) were used at a concentration of 10 µg/ml in 3% BSA/PBS and incubated for 90 minutes. Subsequently cells were rinsed with PBS and washed twice for 5 minutes with 0.1% Tween-20 in PBS. After a short rinse with water the coverslips were mounted onto coverslides (Carl Roth, H879.1) with ProLong Glass Antifade Mountant (ThermoFisher, P36980) and cured over night at room temperature.

### Live-cell imaging of COS-7 and U2Os cells

Cos7 or U2OS cells were seeded on Cell Imaging Coverglass, 8 Chambers from Eppendorf (0030742036) and incubated at 37°C and 5% CO_2_. After 24h cells were transfected with FuGENE® HD (Promega, E2311) in a ratio of 4:1 FuGENE (µl) : DNA (µg). Cells were incubated 24h -30h before imaging. Data of EMTB-3xGFP and tdTomato-Calreticulin was acquired simultaneously on a ZEISS Elyra 7 system with Duolink (two 4.2 CL HS pco.edge sCMOS cameras) using the 488 nm and 561 nm laser lines, Plan-Apochromat 63x 1.4 oil objective, 13 phases, 8 ms exposure time and 37°C. Simultaneous imaging allowed burst mode processing. mEmerald-Rab5a in U2OS cells was imaged with 9phases and 1 ms exposure time at 37°C.

### Plasmids

The EMTB-3XGFP plasmid was a gift from William Bement (Addgene plasmid # 26741; http://n2t.net/addgene:26741 ; RRID:Addgene_26741) to the University of Wuerzburg.^[61]^ The tdTomato-Calreticulin-N-16 and mEmerald-Rab5a-7 plasmids were a direct gift from Michael Davidson to Zeiss and are also documented on Addgene (Addgene plasmids 58074 and 54243).

### Immunolabeling of gp210 in nuclear envelopes

*Xenopus laevis* oocyte nuclear envelopes were prepared exactly as described in Loeschberger *et al*. with two exceptions.^[46]^ For the isolation of oocyte nuclei and nuclear envelopes “5:1 medium” (83 mM KCl, 17 mM NaCl, 1.5 mM KH_2_PO_4_, 7 mM Na_2_HPO_4_) was used. Nuclear envelopes attached to coverslips were fixed for 20 min with 2% paraformaldehyde in “5:1 medium”. Envelopes were saturated in 0.5% BSA/PBS for 10min, incubated with primary antibody against gp210 (1:300 in PBS) for 45 min, washed in PBS and incubated 30 min with alpaca anti-mouse Alexa Fluor 488 secondary antibody (1:300 in PBS). The central channel of NPCs was labeled with 1µg/ml Alexa Fluor 555 labeled wheat germ agglutinin (WGA) in PBS for 10 min. Envelopes were embedded in ProLong Glass Antifade mountant, cured overnight and stored at 4°C until imaging. The mouse monoclonal antibody ASC222a2 against gp210 was provided by Georg Krohne.^[62]^ Alpaca anti-mouse IgG1 coupled to Alexa Fluor 488 was purchased from Chromotek (sms1AF488-1-100).

### Mouse brain tissue sample

The Thy1-GFP mouse brain sample was a kind gift from Jochen Herms to Zeiss and is described in detail in literature.^[63,64]^

### Lattice-SIM and SIM-Apotome of fixed samples

SIM imaging was performed on a ZEISS Elyra 7 system, using 488, 561 and 641 nm laser lines. For SIM-Apotome imaging of the Thy1 brain tissue sample we used the LCI Plan-Apochromat 25x/0.8 multi immersion and the Plan-Apochromat 40x/1.4 oil objective. Five raw images (phases) per final SIM-Apotome image were aquired. Lattice SIM imaging needed 13 phases and was generally performed with the Plan-Apochromat 63x 1.4 oil objective, with two exceptions: For the images in Figure S3 (Supporting Information) the alpha Plan-Apochromat 63x/1.46 oil corr and for the images of the nuclear envelope in Figure S5 (Supporting Information) the alpha Plan-Apochromat 100x/1.57 Oil-HI was used.

### Resolution verification with technical samples

Resolution of the *di*SIM processing was verified using a customized 60 nm DNA Nanoruler with Alexa Fluor 488 (GATTAquant GmbH) and the Argo-SIM slide (ARGOLIGHT SA, Argo-SIM SLG-800) using the 488 nm laser line.

### Estimating the lateral resolution in biological samples with Fourier Ring Correlation

To estimate the lateral resolution in complex biological samples we employed FRC analysis.^[48,65,66]^ Data were analyzed using the FIJI-plugin by O. Burri and A. Herbert (https://imagej.net/Fourier_Ring_Correlation_Plugin). Providing two statistically independent data sets of the same sample, we first performed the SIM/*di*SIM reconstruction and then analyzed individual z-planes with FRC. Although the unprocessed camera images often exhibited unwanted correlations at high frequencies -presumably due to the fixed pattern noise of the sCMOS camera present in both data sets and thus destroying the statistical independence -the SIM/*di*SIM reconstruction seemed more robust due to multiple steps in the processing pipeline where information is mixed from various pixels.

### *di*SIM *processing*

All *di*SIM processing was performed in ZEN Black (Figure 1). The *di*SIM processing (Figure 1g-1m) differs in two central aspects from the classical linear SIM processing (Figure 1a-1f). First, the deconvolution part is separated from order combination, thus allowing out-of-focus suppression and a reduction of the noise due to the summation of the various images before entering the iterative deconvolution. Second, all processing effects especially the blocking of out of focus light by Fourier filtering the zero order and the suppression of the grating frequencies in higher orders will be reflected in the PSF. Failure to do so usually leads to divergence of the iterative deconvolution limiting the number of possible iterations to 3-5 at maximum^[25]^ and potential artefacts in the images.

In SIM, the suppression of the background arising from fluorescence from outer focal areas is so successful, since the suppressed frequencies can be restored from other orders. For the same reason SIM even allows an initial smoothing of the raw data reducing the high frequency noise in the images.^[24]^ Thus, the data and PSF prepared in the same way are extremely well suited to the nonlinear DCV methods which only achieve their full potential by an accurate PSF and high contrast data.

The successfully applied weighting of the orders in the classical SIM reconstruction (Figure 1e) which boosts the high frequencies and ensures that this information is not drowned in the noise of the other orders when summed for combination could also potentially be applied in *di*SIM. However, the weighting as in the classical processing requires a Wiener filter to deal with the OTF approaching zero at the edges. Using the same approach for *di*SIM would introduce Wiener artifacts propagating into the iterative deconvolution. In addition, for a successful iterative deconvolution the noise levels need to be kept low. This is more critical than in the classical reconstruction. The boost of the higher frequency orders will also lead in some cases to an increase of the noise, which in many cases is counterproductive for the subsequent iterative deconvolution. Thus, we applied no OTF weighting so far. Adapting the classical OTF weighting to the *di*SIM noise sensitivity might improve the method further.

## Acknowledgements

The authors thank G. Krohne from the Department of Electron Microscopy, Biocenter, University of Würzburg for the preparation of oocyte nuclear envelopes. We thank Dr. Kai Wicker from Corporate Research and Technology of Carl Zeiss AG for carefully reading of the manuscript and valuable comments. T.K. and M.S. acknowledge support by the German Research Foundation (DFG SA829/19-1 and TRR 166 ReceptorLight, project A04) and the European Regional Development Fund (EFRE project: “Center for Personalized Molecular Immunotherapy”).

## Supporting Information

**Figure S1.**
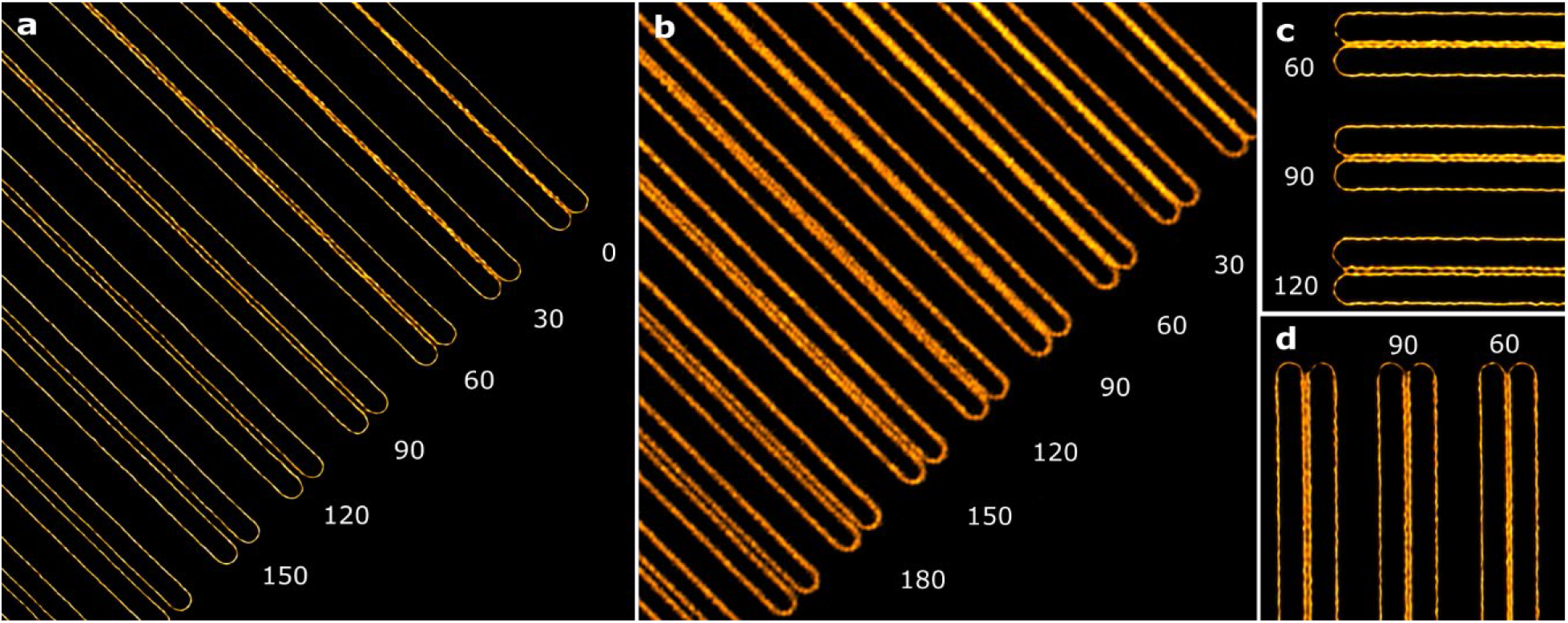
Estimating the spatial resolution of *di*SIM with Argo-SIM slides. Z-stacks with Lattice-SIM acquisition were taken at 488 nm excitation and 500-550 nm detection bandwidth. Values are given in nanometers. a, b) Comparison of the 3D SIM resolution targets resolved by *di*SIM (a) and generalized Wiener reconstruction (b). c-d) Horizontal and vertical direction of the same resolution target resolved by *di*SIM.

**Figure S2.**
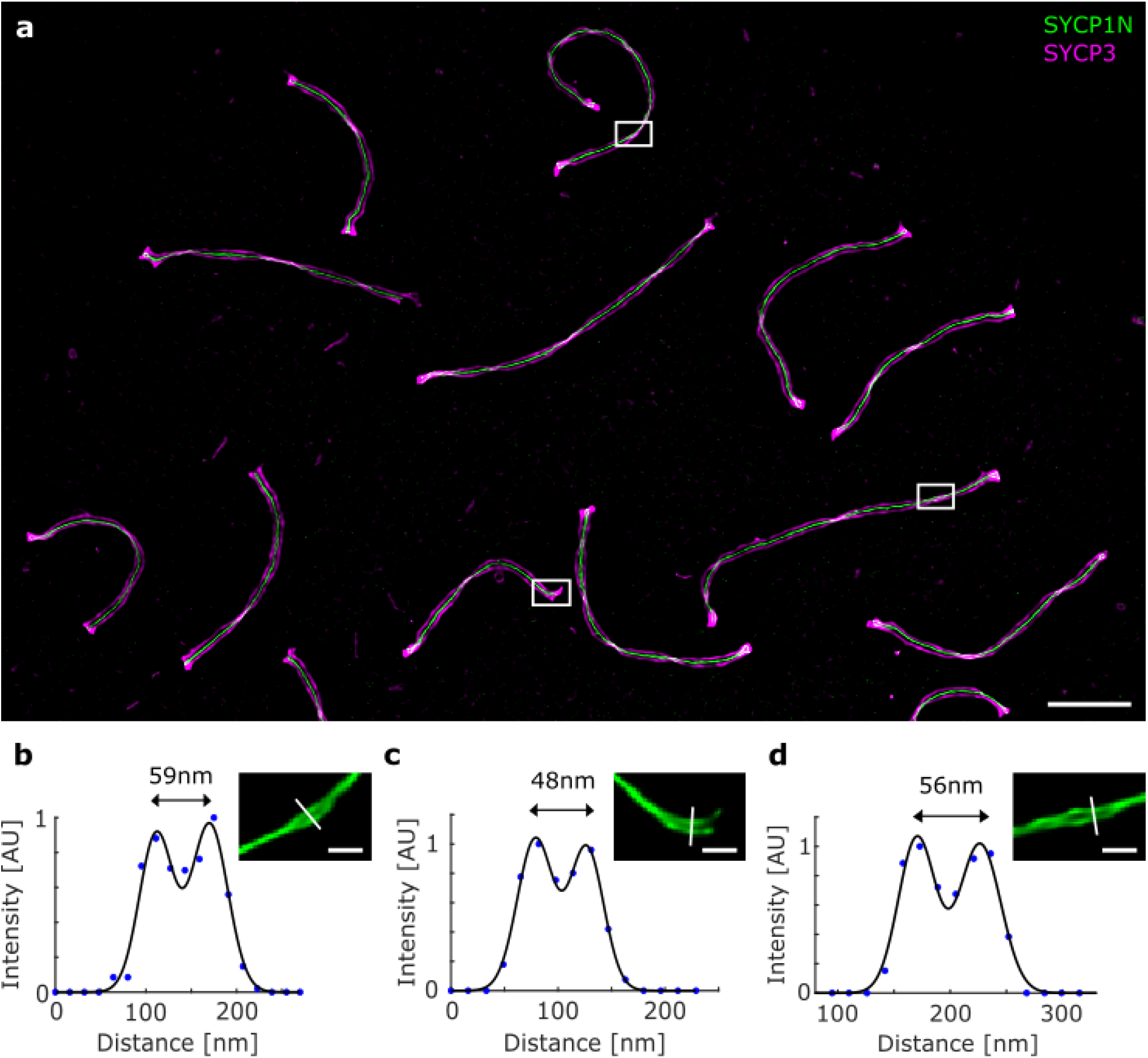
*di*SIM resolves the molecular organization of SCs with 50-60 nm spatial resolution. a) Two-color *di*SIM image of SC spreadings. The transverse filament and lateral element proteins SYCP1-N and SYCP3 were labeled with Alexa Fluor 488 (green) and Alexa Fluor 647 (magenta), respectively. b-d) In lateral view sections, where the SYCP3 signal twists, we determined the distances between the SYCP1-N signals for three selected regions close to the twisting points to 48-59 nm to demonstrate the superior resolution of *di*SIM. Recently, standard SIM images of post-expansion immunolabeled SCs supported a multilayered organization of SYCP1-N in lateral views.^[1]^ The occasional disclosure of multimodal distributions of the SYCP1-N signal in some lateral view sections of expanded SCs was attributed to the higher labeling density enabeled by post-expansion immunolabeling. Scale bars, a) 2.5 µm, b-d) 250 nm

**Figure S3.**
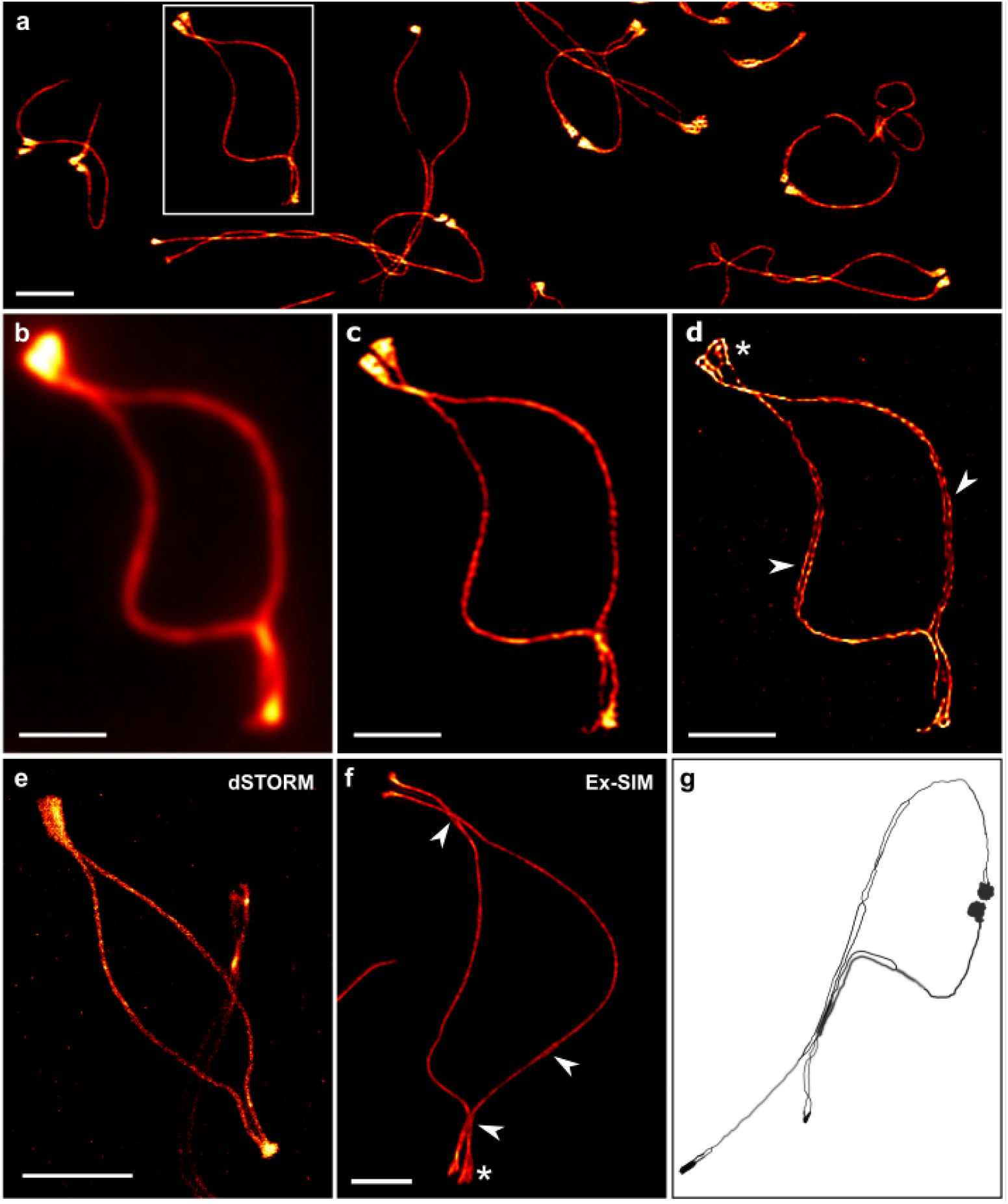
Molecular details of the lateral elements of the SC resolved by *diSIM*. a) SIM of murine spermatocyte spreadings. SYCP3 (red) serves as a molecular marker for the lateral element of the synaptonemal complex (SC). The boxed region in (a) shows the SYCP3 signal associated with a bivalent at the late prophase stage of diplonema. b-d) Different reconstructions of the boxed region highlighted in (a). b) Wide-field fluorescence, c) conventional SIM, and d) *di*SIM image. Since *di*SIM provides improved lateral resolution, the molecular architecture of the SC such as the fraying of the SYCP3 signal (d asterisk, arrowheads) can be elucidated in detail. e) *d*STORM cannot resolve the bifurcation (arrowheads) and fraying (asterisks) of the SYCP3 signal characteristic for later prophase stages such as diplonema. f) The fine structural details shown with *di*SIM here were only recently resolved by expansion microscopy combined with SIM (Ex-SIM).^[1]^ g) Previously, the morphological equivalent was solely observed in EM micrographs, based on which a role of the two sister chromatids in the compaction of multiple SYCP3 positive strands into two sub-lateral elements that fray at the ends was proposed (schematic representation adapted from literature.^[2]^ Scale bars, a) 2 µm, b-f) 2 µm,

**Figure S4.**
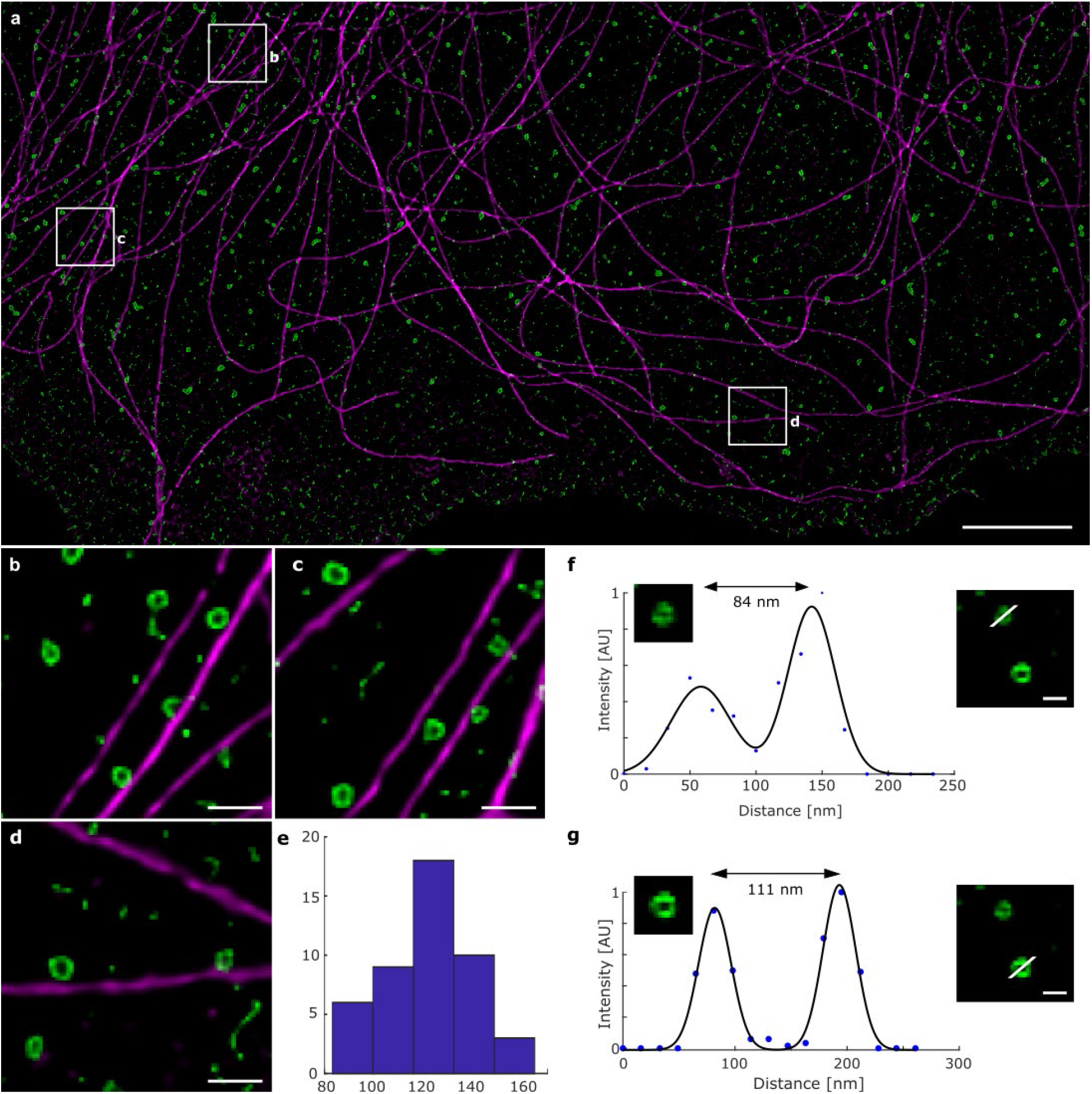
*di*SIM of clathrin-coated pits as cellular reference structure. a) *di*SIM image of an immunostained COS-7 cell labeled with anti-alpha-tubulin primary and Alexa Fluor 568 secondary antibodies (magenta), and anti-clathrin primary and Alexa Fluor 488 secondary antibodies (green). For better visibility of CCPs the gamma of the green and magenta channel were adjusted to 0.7 and 0.9, respectively, in ZEN black. b-d) Expanded views of the boxed regions in (a). e) Fitting of the histogram of diameters of 45 CCPs results in an average diameter of 122.4 ± 19.5 nm (s.d.). f,g) Diameters of CCPs were determined with Gaussian fits to line profiles across the pits. Here two examples of CCPs with diameter of 84 nm (upper graph) and 111 nm (lower graph) are shown. Scale bars, a) 5 µm, b-d) 500 nm, f,g) 200 nm.

**Figure S5.**
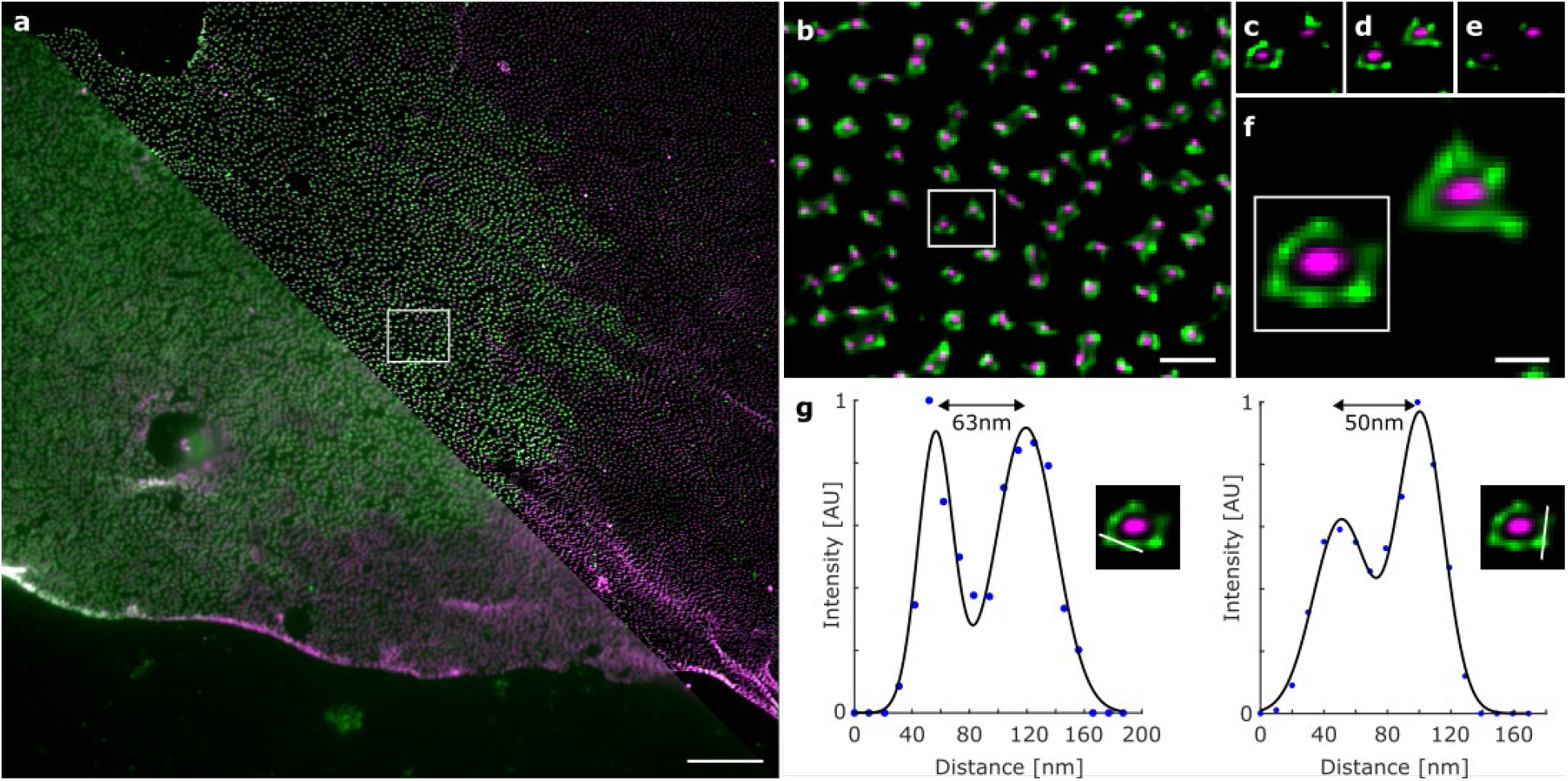
NPCs show the limits of *di*SIM. a) Two-color fluorescence images of a nuclear envelope from a *Xenopus laevis* oocyte recorded by wide-field (lower left part) and linear SIM (upper right part). The outer ring of nuclear pore complexes (NPCs) was immunolabeled with anti-gp210 primary and Alexa Fluor 488 secondary antibody (green). The central channel was labeled with Alexa Fluor 555 labeled wheat germ agglutinin (WGA) (magenta). b) Expanded *di*SIM image of the white rectangle in (a). c-e) Different z-planes of two NPCs indicated by the white rectangle in (b) are depicted to demonstrate the sectioning capabilities of *di*SIM. f) Maximum intensity projection image of the two NPCs for better visualization of the ring structure. g) *di*SIM resolves subunits of the outer ring structure of NPCs. For a small population of not too densely packed NPCs *di*SIM resolves individual gp210 subunits separated in the two examples shown by 63 nm and 50 nm. The central channel of NPCs with a diameter of ∼40 nm cannot be resolved by *di*SIM. Scale bars, a) 5 µm, b) 500 nm, f) 100 nm.

**Figure S6.**
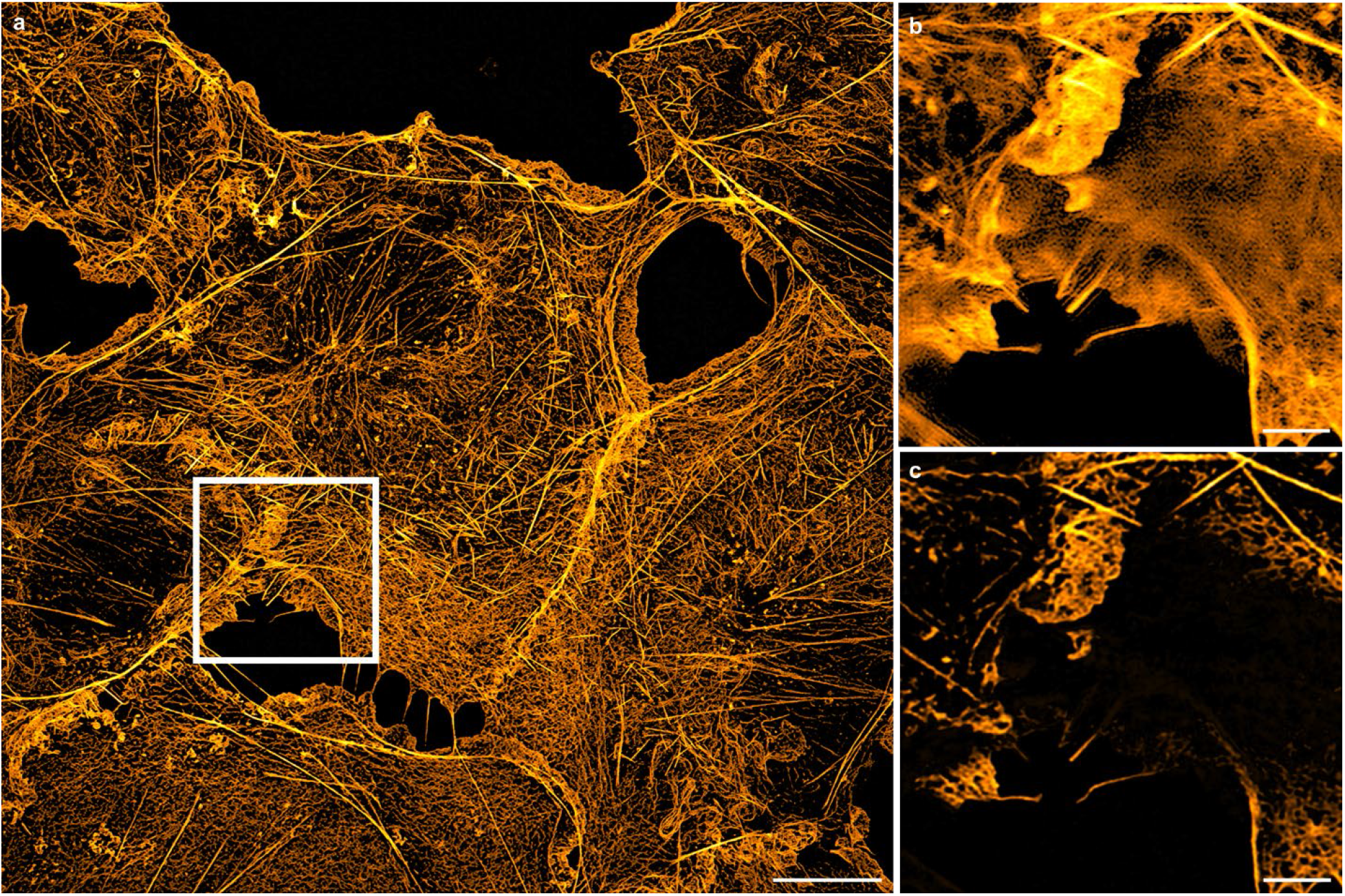
SIM artifacts are effectively removed by *di*SIM processing. Actin cytoskeleton stained with Alexa Fluor 488-phalloidin in COS-7 cells. a) Maximum intensity projection of a *di*SIM reconstruction imaged with Lattice-SIM. b,c) Single z-planes of the white rectangle marked in (a) to compare a conventional Wiener filter reconstruction exhibiting a strong meshwork of structured noise and out-of-focus blur overlaying the actin structures (b) with *di*SIM reconstruction (c). To better visualize the otherwise dim and small filamentous structures a display gamma of 0.45 was used. Scale bars, a) 10 µm, b,c) 2.5 µm.

**Figure S7.**
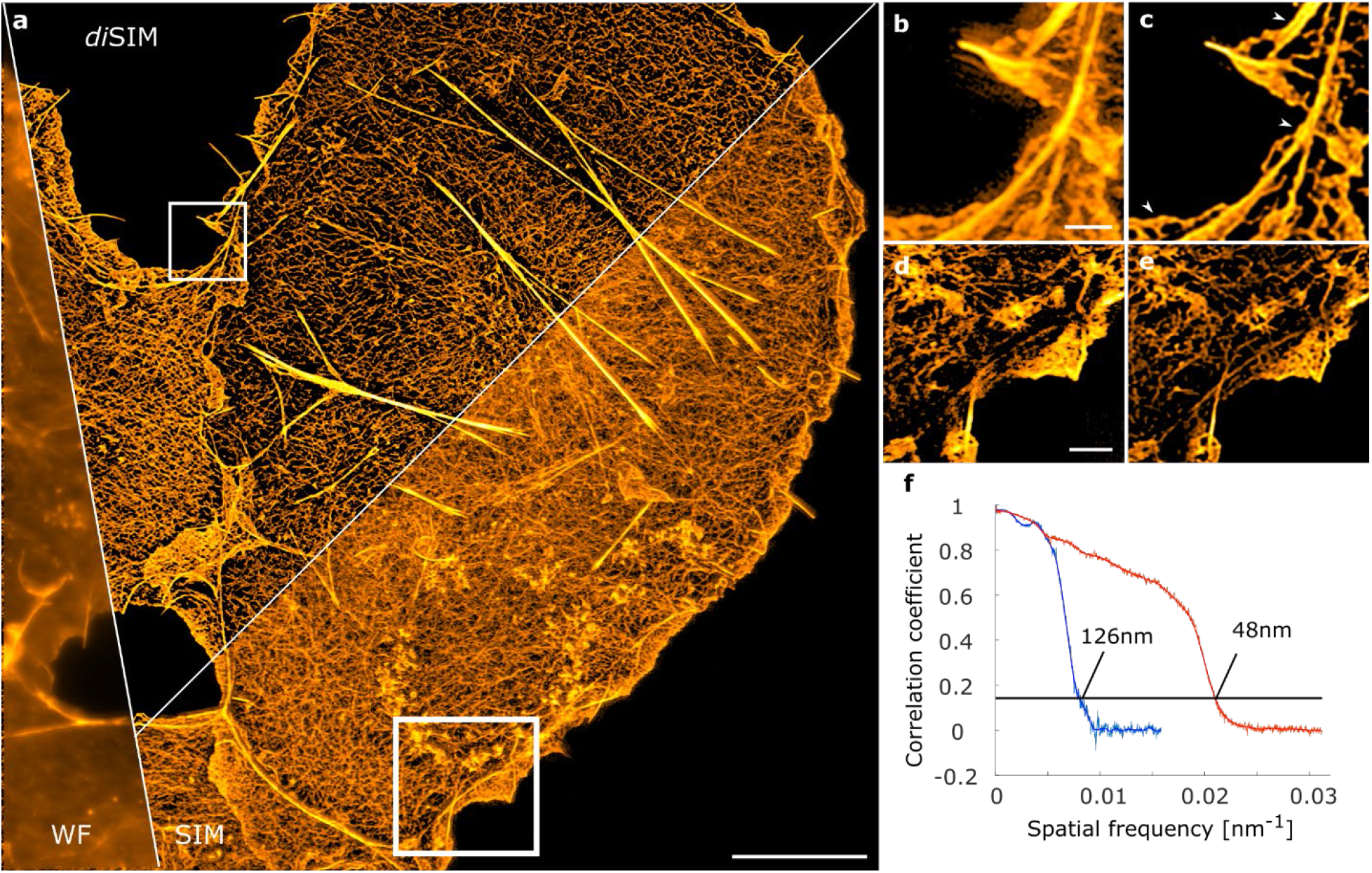
Enhanced resolution in *di*SIM reconstructions assessed by Fourier Ring Correlation analysis. Actin cytoskeleton stained with Alexa Fluor 488-phalloidin in COS-7 cells. a) Maximum intensity projections of wide-field (WF), conventional SIM and *di*SIM reconstructions imaged with Lattice-SIM illumination. b,c) Comparison of expanded SIM (left) and *di*SIM (right) reconstructions of the white rectangle in the *di*SIM image of (a) illustrate not only improved sectioning and artifact suppression but especially increased spatial resolution (white arrows). d,e) Expanded SIM (d) and *di*SIM (e) reconstructions of the white rectangle in the SIM image of (a). (**f**) Fourier ring correlation (FRC) analysis^[3-5]^ of two independent datasets of the same sample illustrating the improved resolution of the *di*SIM reconstruction (red) as compared to the standard SIM reconstruction (blue). The conventional resolution cutoff (1/7) is indicated by the black line with the corresponding maximal resolution displayed in the blue and red curve. Scale bars, a) 10 µm, b,c) 1 µm, d,e) 2 µm.

**Table S1.** Primary antibodies used for the immunolocalization of SC proteins on nuclear spreadings.

| Primary Antibody | Epitope | Dilution |
| --- | --- | --- |
| Mouse anti-SYCP3 (ab97672; Abcam) | full length | 1:100 |
| Guinea-pig anti-SYCP1 N-terminus (54) | aa 1-124 | 1:100 |
| Rabbit anti-SYCP1 C-terminus (55) | aa 922-997 | 1:150 |
| Rabbit anti-SYCE3 (56) | full length | 1:150 |

**Figure S9.**
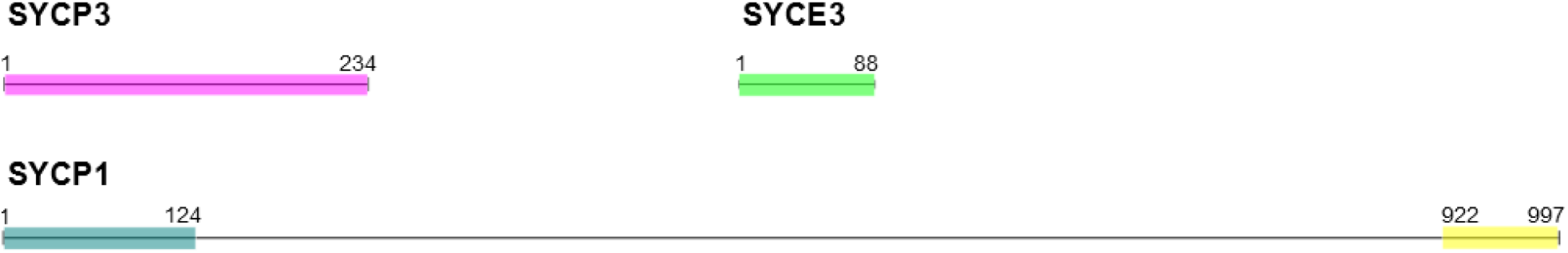
Schematic representation of the full length protein sequences of SC proteins SYCP3, and SYCP1. Epitopes of these proteins that were targeted by immunofluorescence staining are highlighted by colored boxes.

### Captions for movies S1-S3

**Movies S1-S3**. Dynamics of microtubules and ER in COS-7 cells. Simultaneous two-color acquisition of Calreticulin-tdTomato and EMTB-3xGFP at 37°C imaged at a frame rate of 81.6 Hz (8 ms exposure time) by lattice-SIM with 13 phases.

**Movie S1**. Corresponds to Fig. 4D, Scale bar 2µm.

**Movie S2**. Corresponds to Fig. 4E, Scale bar 1µm.

**Movie S3**. Corresponds to Fig. 4F, Scale bar 1µm.

